# Distinct neural systems for the production of communicative vocal behaviors

**DOI:** 10.1101/107441

**Authors:** ZK Agnew, L. Ward, C. McGettigan, O. Josephs, SK. Scott

## Abstract

Vocalizations are the production of sounds by the coordinated activity of up to eighty respiratory and laryngeal muscles. Whilst voiced acts, modified by the upper vocal tract (tongue, jaw, lip and palate) are central to the production of human speech, they are also central to the production of emotional vocalizations such as sounds of disgust, anger, laughter and crying. Evidence suggests that the speech and emotional vocalizations may comprise distinct classes of vocal movements: patients with speech production deficits are often able to produce emotional vocalizations. In addition to this, the ontogeny of the two articulations is largely distinct, with some culturally universal emotional vocalizations emerging soon after birth and human speech being a culture specific, highly learnt motor act, which must develop to some degree before a critical period in development. Here we investigate the motor pathways underlying these two classes of vocal movements for the first time by directly comparing BOLD responses during production of speech and non-speech vocal movements. Using functional magnetic resonance imaging, we report distinct patterns of activity in both subcortical and cortical regions (putamen and bilateral inferior frontal and parietal cortices) during the production of emotional vocalizations compared to speech production. In contrast we show that responses in primary sensorimotor regions do not differ during the production of speech and emotional vocalizations, suggesting partially overlapping, and partially non-overlapping neural structures for the motor control of these two classes of movement. In addition to this we report that responses in auditory cortices are distinct during the production of speech and non-speech vocalizations, suggesting that feedback control of speech and emotional vocalizations are distinct. These data provide novel evidence for the presence of dual pathways for the neural control of complex articulatory movements in humans. These findings are discussed in relation to the clinical and primate literature of vocal motor control.

**Significance Statement:** This work marks the first evidence in healthy humans for dual routes to vocal behaviors. Clinical evidence suggests that patients unable to produce speech may still be able to produce other vocal behaviors that employ the same set of effectors. Here we demonstrate that the production of different classes of vocal behavior is associated with overlapping and distinct networks of activity. Moreover, we show that auditory processing that occurs during the production of these movements may be distinct, suggesting that feedback may be used differently for these distinct classes of vocal movement.

## Introduction

Vocalizations are the production of a sound by the coordinated activity of respiratory and laryngeal muscles. Whilst vocalizations, modified by the upper vocal tract (tongue, jaw, lip and palate) are central to the production of human speech, they are also central to the production of emotional vocalizations such as sounds of disgust, anger, laughter, crying and so forth (Jurgens, 2002). Speech is only one of many movements made by the human upper vocal tract, yet it is by far the most well-described of the articulatory movements. Whilst recent years have seen a rapid increasing interest in the understanding of nonverbal behaviors, such as emotional vocalizations, gesture and body language, the neural processing underlying these important forms of communication remain largely unknown (Buck & VanLear, 2002).

A number of lines of evidence suggest that the neural control of speech production may be fundamentally different to that of other non-speech sounds such as emotional vocalizations, the latter sharing semblance with vocalizations made by non-human primates than to human speech. The ontogeny of human emotional vocalizations is very different to that of human speech, in fact sharing semblances with the development of non-human primate calls. Human infants, as well as ape and monkey infants display spontaneous vocalizations from early in development; human cries can occur minutes after birth, with laughter appearing in the following weeks. In addition to this, evidence suggests that the acquisition and maintenance of these vocal movements is distinct.

## The role of feedback in different vocal motor behaviors

Auditory feedback is critical for speech acquisition in development (Osberger & McGarr, 1982; Smith, 1975). Additionally, speech production of post lingual deaf adults is selectively altered (Cowie, Douglas-Cowie, & Kerr, 1982), suggesting that speech is responsive to, yet not wholly dependent on feedback, once speech has been learnt (Cowie et al., 1982; Lane & Webster, 1991). The role of auditory feedback appears to be very different for emotional vocalizations. Laughs and cries occur in human infants with loss of sight and hearing (Eibl-Eibesfeldt, 1970). The emotional vocalizations produced by blind and deaf children do not appear to be acoustically altered by the absence of auditory input (Makagon, Funayama, & Owren, 2008) and appear at the same point in development (Scheiner, Hammerschmidt, Jurgens, & Zwirner, 2002, Scheiner, Hammerschmidt, Jurgens, & Zwirner 2004). Finally, unlike speech, many emotional vocalizations are recognized across different cultures (Sauter, Eisner, Ekman, & Scott, 2010). Owren et al., (2011) suggest two principles that may serve to distinguish between different types of primate vocal behavior. The first principle differentiates between the role of auditory-motor experiences of ‘production-first’ (such as emotional vocalizations) and ‘reception-first’ (such as speech) during vocal development. The second principle refers to evidence suggesting distinct neural pathways controlling different vocal behaviors, which we investigate in the present study.

## Potentially distinct neural pathways for speech and emotional vocalizations

Owrens and colleagues (2011) posit that distinct neural systems for different vocal behaviors are a second important defining feature of vocal behavior. They suggest that the evolution of a new vocal behavior in vocal primates does so via the development of a second neural pathway, as opposed to an existing network for vocal behavior, suggesting that emotional vocalizations may be served by distinct neural systems from those underlying speech. Second, clinical data from a number of different patient groups indicates at last a partial dissociation between the neural systems underlying the production of speech and emotional vocalizations. Clinically, patients with speech apraxia have impaired speech production, but they spontaneous production of emotional vocalizations (e.g. laughter) often remains intact. Whilst this may originate from neural differences in the spontaneous production of action and cued action, others have reported similar findings for cued speech and emotional vocalizations. For example, Rohrer, Warren and Rossor (2009) report a group of ten patients with primary progressive aphasia leading to mutism. In these patients, progressive loss of speech output was accompanies by abnormal laugh like vocalizations, which they report did not retain contextual sensitivity but instead ‘replaced speech’. These data indicate that following the disruption of the fronto-temporal networks underlying speech production; patients may have resorted to using intact neural networks subserving laughter production to attempt to make some form of vocalization. Similarly, Cummings et al. (1983) report that it is not unusual for patient with mutism to retain some phonatory ability in emotional utterances, which may be evidence of an ability to utilize subcortical systems for communication in the face of cortical damage to speech output systems (Larson, 1988). Given these differences, it has been suggested that emotional vocalizations may rely on evolutionarily older, or different, (potentially/partially affective and subcortical) neural systems.

In sum, significant evidence exists to suggest that human emotional vocalizations may fall under at least partially distinct neural control to that of human speech. At present, no study has looked at the neural basis of emotional vocalization production with respect to that of speech production. Here we aim to investigate how the neural basis of these classes of vocal behaviors (speech and emotional vocalizations) differs, both in terms of motor output and sensory reafference.

## Methods

### Subjects

Fifteen right-handed speakers of English (6 female, mean age 26) took part in the present study. All participants had normal hearing and no history of speech/ language, neurological or psychiatric problems (selfreported). All subjects were naive to the purpose of the experiment. This study was approved by the University College London Department of Psychology Ethics Committee.

### Conditions

The experimental set up required the following seven conditions, conditions 5 and 6 are not included in this manuscript:

1. Speech production (SpProd)
2. Emotional vocalization production (EmoVocProd)
3. Silent speech production (SpSilent)
4. Silent emotional vocalization (EmoVocSilent)
5. Covert speech production (SpCov)
6. Covert emotional vocalization (EmoVocCov)
7. Silent rest (Rest)

During the main experimental run, subjects lay supine in the scanner in the dark and were asked to pay attention to instructions presented visually. The instructions were to get ready to either to produce words, produce emotional vocalizations, to silently produce or mime words or emotional vocalizations related to a given emotional category, or to imagine producing words or emotional sounds. The four emotional categories were Amusement, Relief, Disgust and Sadness, two positively and two negatively valenced (Ekman, 1992, 2003), all considered to be culturally universal (Sauter et al., 2010). In the speech production condition, subjects were trained to produce words related to that condition. For example, following the cue ‘Get ready to produce speech related to *AMUSEMENT*’, a fixation cross would appear during which time subjects would produce words such as ‘happy’, ‘smile, ‘laugh’, ‘fun’ and so on. In the emotional vocalization condition, subjects were trained to produce sounds that would be appropriate if they were feeling that emotion. They were instructed not to try and ‘act’ but to think about the sorts of sounds they would genuinely produce when feeling that emotion, even if the vocalization did not produce a great deal of sound. Subjects were trained sufficiently that they did not find it hard to generate appropriate words and sounds. Following the instruction to get ready subjects were presented with a fixation cross for 3 seconds, during which they were to perform the task. This provided sufficient time to produce 3 utterances. Subjects were instructed to stop when the fixation cross disappeared from the screen. Instructions were presented using matlab (Mathworks, version 7.10) using the Psychophysics Toolbox extension (Brainard, 1997).

### Auditory and Motor Localizer

A independent localizer (continuous acquisition, blocked design) was run in the same subjects, to identify cortical regions active during movements made at the lower end of the vocal tract (glottal stop), at the upper end (lip pursing) facial movements (alternating smile and nose wrinkle), and passive listening to pseudorandomised emotional vocalisations and speech. The peaks from these clusters identified by these motor conditions were used to make 6mm diameter spherical regions of interest from which the beta values were extracted.

### Functional MRI

Functional imaging data were acquired on a Siemens Avanto 1.5-T scanner (Siemens AG, Erlangen, Germany). This type of head coil has been shown to increase signal-to-noise for medical images acquired in the 1.5 Tesla field without increasing the signal drop out associated with higher field strengths (Fellner et al., 2009; Parikh et al., 2011). A single run comprising 370 echo-planar whole-brain volumes was acquired, with a 2mm slice thickness, TA= 3700ms, TR =16.1s, echo time = 24.26 msec, flip angle = 90°, 40 axial slices, 3 mm × 3 mm × 3 mm in-plane resolution). A clustered sparse-sampling routine was employed (Edmister, Talavage, Ledden, & Weisskoff, 1999; Hall et al., 1999), in which three volumes were acquired per trial and whereby the task was performed in a 5 second silent period between clusters of scans. This approach serves to reduce head motion related artifact associated with vocalization by ensuring that data acquisition begins after then offset of vocalization.

The experiment comprised of 6 sessions, with 20 trials per session, organized such that three volumes were acquired per trial, comprising 360 volumes. Including dummy scans the total number of scans acquired was 370. There were 120 trials in total. The entire experiment lasted just over 30 minutes (7 conditions x 120 trials x 16.1sTR).

During each run, subjects were cued to perform one of each of the six experimental conditions or to rest silently. There were 17 examples of each condition, the order of which was pseudo-randomized to allow a relatively even distribution of the seven conditions in the absence of order effects. The first two functional volumes were discarded in order to remove the effect of T_1_ equilibration. High-resolution T_1_ anatomical volume images (160 sagittal slices, voxel size 1mm^3^) were also acquired for each subject.

### Stimulus presentation

Stimuli (visually presented prompts) were presented using MATLAB with the Psychophysics Toolbox extension. Instructions were projected from a specially-configured video projector (Eiki International, Inc., Rancho Santa Margarita, CA) onto a custom-built front screen, which the participant viewed via a mirror placed on the head coil. Speech output was recorded using Audacity (http://audacity.sourceforge.net/). The audio channel was routed through a Sony HD-510 amplifier (Sony Europe Limited, Weybridge, UK) to electrodynamic MR-compatible headphones worn by the participant (Sensimetrics Corporation, Malden, MA). Instructions were presented via front-projection from an EIKI LC-XG250 projector (Eiki International, Inc., Rancho Santa Margarita, CA) to a custom-built screen at the mouth of the scanner bore, which was viewed using a mirror placed on the head coil.

Pre-processing and analyses: Functional data were analyzed using SPM8 (Wellcome Department of Imaging Neuroscience, London, UK) running on matlab 7.4 (Mathworks Inc, Sherborn, MA). All functional images were realigned to the first volume by six-parameter rigid body spatial transformation. Functional images were normalized into standard space using the Montreal Neurological Institute (MNI) template using parameters elicited from a unified segmentation of the T1 anatomical image. Functional images were then smoothed using a Gaussian kernel of full-width half-medium (FWHM) 8 mm. Event-related responses for each condition were modeled as a finite impulse response function. Event onsets were modeled from the onset of auditory recording with a 4 second duration. Each condition was modeled as a separate regressor in a general linear model. Motion parameters (three translations and three rotations) were modeled as six regressors of no interest at the single-subject level. For each subject (first level), contrast images were created to describe the comparisons between each of the experimental conditions compared to silent rest and to each other. The contrast images generated from these t-tests were entered into group analyses at the second level. Peak activations were localized using the anatomy toolbox available within SPM8 ( Eickhoff et al., 2005). In all cases statistical maps were thresholded at p<0.001 with a cluster extent of 30 voxels.

### Region of interest analyses

Region of interest analyses were carried out to investigate mean effect sizes in specific regions across all experimental conditions against baseline, using the MarsBar toolbox that is available for use within SPM8 (Brett et al., 2002). Regions of interest were selected post-hoc, using peaks from contrasts of interest to investigate the profile of activity in these regions across other conditions. Statistical comparisons were not applied to the extracted effect sizes so as to avoid ‘double dipping’ (Kriegeskorte, Simmons, Bellgowan, & Baker, 2009). Second-level clusters were used to extract condition-specific parameter estimates from regions of interest (ROIs) (using MarsBaR Brett, Anton, Valabregue, & Poline, 2002). The anatomical locations of peak and sub-peak voxels (at least 8mm apart) were labeled using the SPM Anatomy Toolbox (version 20) (Eickhoff et al., 2005).

## Results

### Neural networks associated with human vocalization

Both production of speech and production of emotional vocalizations were associated with similar distributed patterns of activity in bilateral sensorimotor cortices, dorsolateral temporal, inferior frontal cortices and cerebellum (see Figure 1, see Table 1 for coordinates).

**Figure 1.**
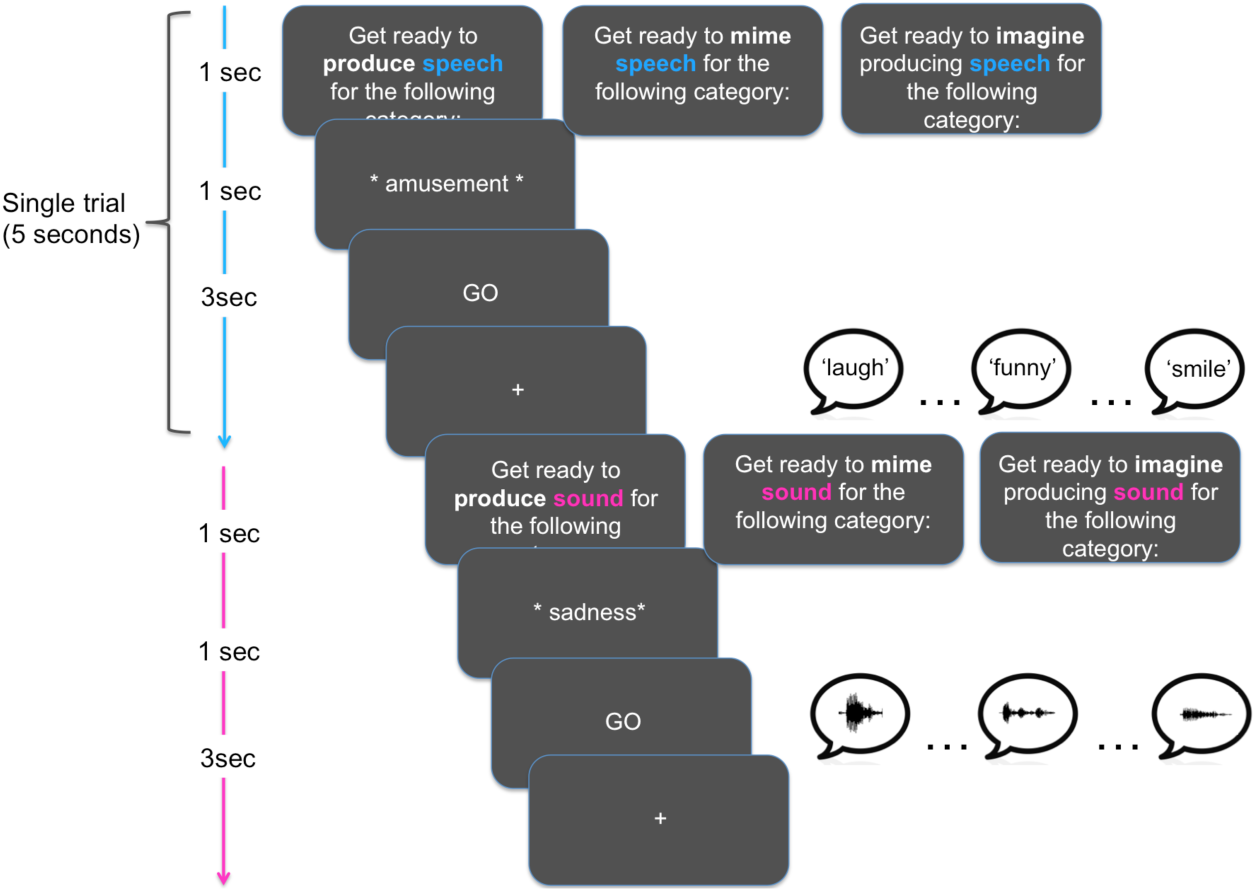
Experimental design across all seven conditions. Subjects underwent extensive training in order to get used to the conditions. In the scanner subjects were cued to produce, mime or imagine producing words, or emotional vocalizations from one of four emotional categories. On each trial they had 3 seconds to produce 3 separate tokens. Sparse imaging was used to minimize the effect of head movement produced as a result of vocalizing. The two covert conditions are not included in this analysis.

## Processing the auditory consequences of speech and emotional vocalizations [Prod > Mime]

By comparing overt production with silent production (miming) we were able to look at activity related to the acoustic processing of ones own articulation in order to investigate whether the auditory response profile differed for the two classes of vocalization. Figure 2 shows maps of activity associated with these comparisons. Production of both speech and emotional vocalizations compared to miming was associated with activity in bilateral STG in a region identified by an independent auditory localizer (white line). The comparison of production with miming for speech and for emotional vocalizations was associated with partially overlapping and partially distinct patterns of activity. There was overlap between the two contrasts in mid STG in both hemispheres, however the two clusters extended in different directions through the temporal cortices. In the case of speech production versus miming, activity in the left hemisphere extended posteriorly and laterally to posterior STG, and then ventrally to the superior temporal sulcus. Outside of auditory sensitive regions, this contrast also revealed clusters of activity in the lingual gyri and cerebellum in both hemispheres, left middle and superior frontal gyri, left superior medial gyrus, left amygdala and right supplementary motor area. The reverse contrast [SpMime > SpProd] revealed a single cluster of activity lying on the border of BA2 of primary somatosensory cortex and the left supramarginal gyrus of the inferior parietal lobe.

**Figure 2.**
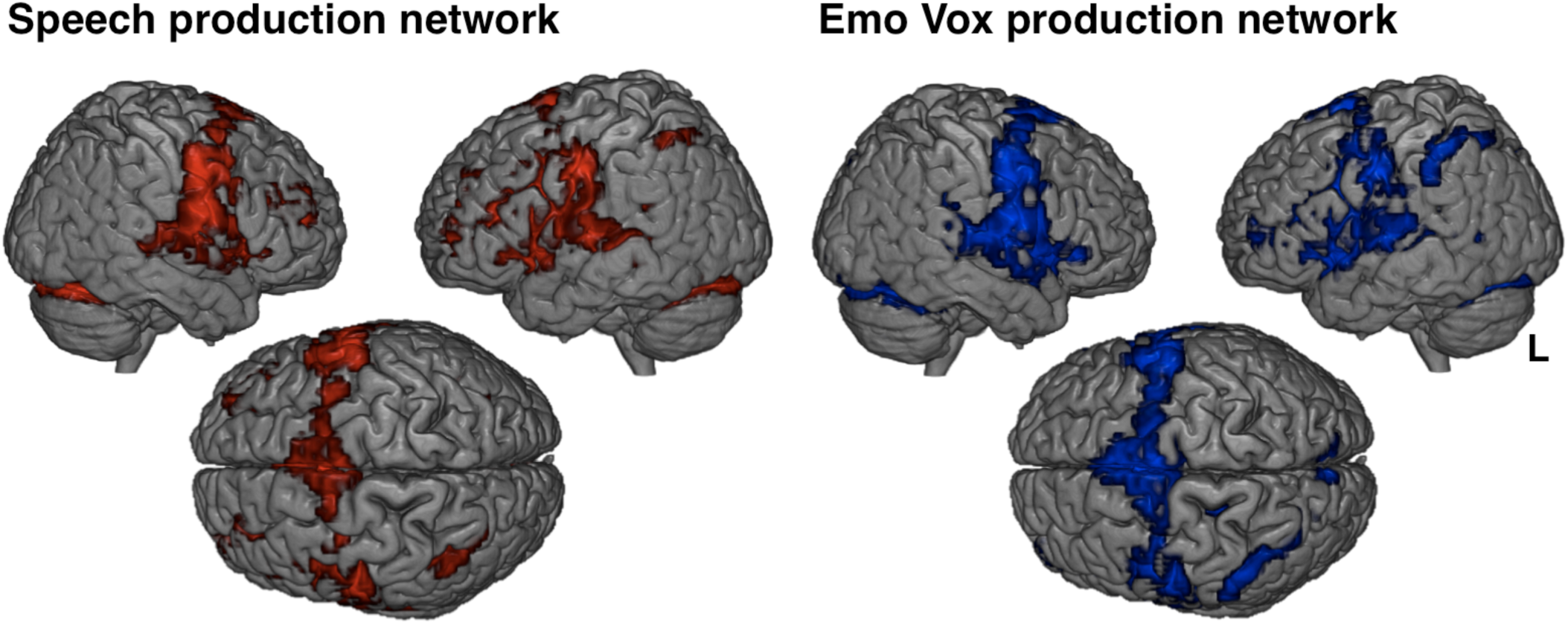
Speech and emotional vocalizations are associated with similar patterns of activity. *Bold responses to production of speech (red) and emotional vocalisations (blue) are seen on a rendered template p<0.001, k=20. BOLD responses during miming of these conditions produced very similar patterns of activity. Direct comparisons are shown in the next section*. Both production of speech and production of emotional vocalizations were associated with similar distributed patterns of activity in bilateral sensorimotor cortices, dorsolateral temporal, inferior frontal cortices and cerebellum.

The same comparison for emotional vocalizations was associated with activity extending from the same mid STG region (overlap between [SpProd>SpMime] and [EmoVocProd>EmoVocMime] is shown in pink) towards anterior STG and into a separate peak in pars opercularis and triangularis of the inferior frontal gyrus. These clusters all lay within the auditory localizer. Outside of these regions, there was also activity in left superior medial gyrus and the right putamen.

Production of emotional vocalizations compared to speech production is associated with increased activity in widespread cortical regions but not primary sensorimotor cortices [EmoVocProd>SpProd] The direct comparison of speech production and production of emotional vocalizations revealed distinct patterns of activity underlying the two articulatory movements. Emotional vocalization production was associated with activity in the right putamen, Crus 1 of the right cerebellum, bilateral regions lying on the boundary between the most ventro-rostral extent of the left supramarginal gyrus where it borders the Sylvian fissue (corresponding to PFop) and OP1 of the rolandic operculum in secondary somatosensory cortex (Eickhoff et al., 2005) left IFG (pars orbitalis, BA47 and pars opercularis, BA44 with a subpeak in pars triangularis, BA45). In the right we report clusters in premotor cortex, middle temporal gyrus, left medial superior parietal cortex (cuneus, area 7P), left insula cortex, and the most ventro-rostral extent of the left supramarginal gyrus where it borders the Sylvian fissue (corresponding to PFop).

There were no whole brain differences in primary motor, somatosensory or auditory cortices. In order to look in further detail at the responses in primary motor cortices a region of interest approach was employed. An independent localizer run in the same subjects was used to identify movements made at the lower end of the vocal tract (glottal stop) and at the upper end (lip pursing). The peaks from these clusters were used to make 6mm diameter spherical regions of interest from which the beta values were extracted. These plots can be seen in Figure 3b. Paired t-tests indicated that there were no significant differences in the betas within these regions during speech production and emotional vocalization production in either hemisphere.

**Figure 3.**
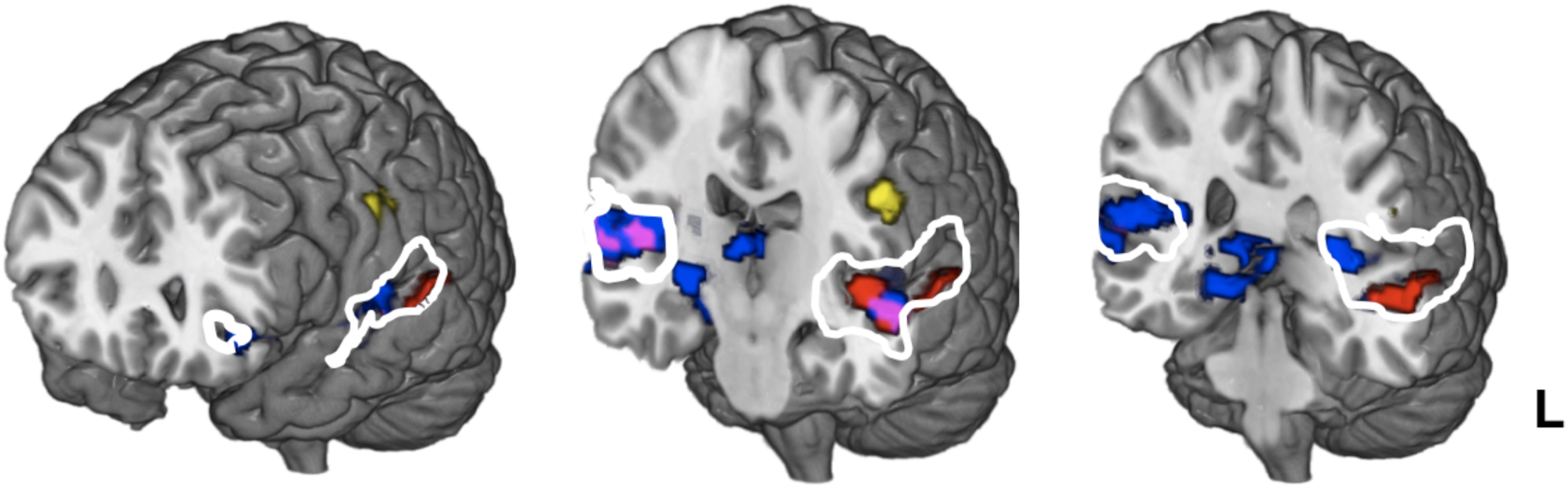
Distinct spatial patterns for mimed speech and emotional vocalizations. The comparison of production with miming allows us to delineate activity associated with hearing the auditory consequences of ones own vocalization. For speech this comparison revealed significant activity in bilateral medial STG extending from a point in mid STG extending posteriorly to the lateral surface (red, [SpProd > SpMime]). The reverse contrast [SpMime > SpProd] was associated with activity left PoCG (area 3b, yellow). The same comparison for emotional vocalizations (blue, [EmoVocProd > EmoVocMime]) also revealed significant activity in mid STG, in an overlapping region (pink), that lies within auditory cortex (defined with an independent auditory localizer and depicted by a white line). This cluster however did not extent in a posterior direction but extended anteriorly and medially, and in the left eventually encompassed the inferior frontal gyrus which also lay within the auditory localizer. Outside of auditory sensitive regions, this contrast also revealed significant activity subcortically in the basal ganglia.

**Figure 4.**
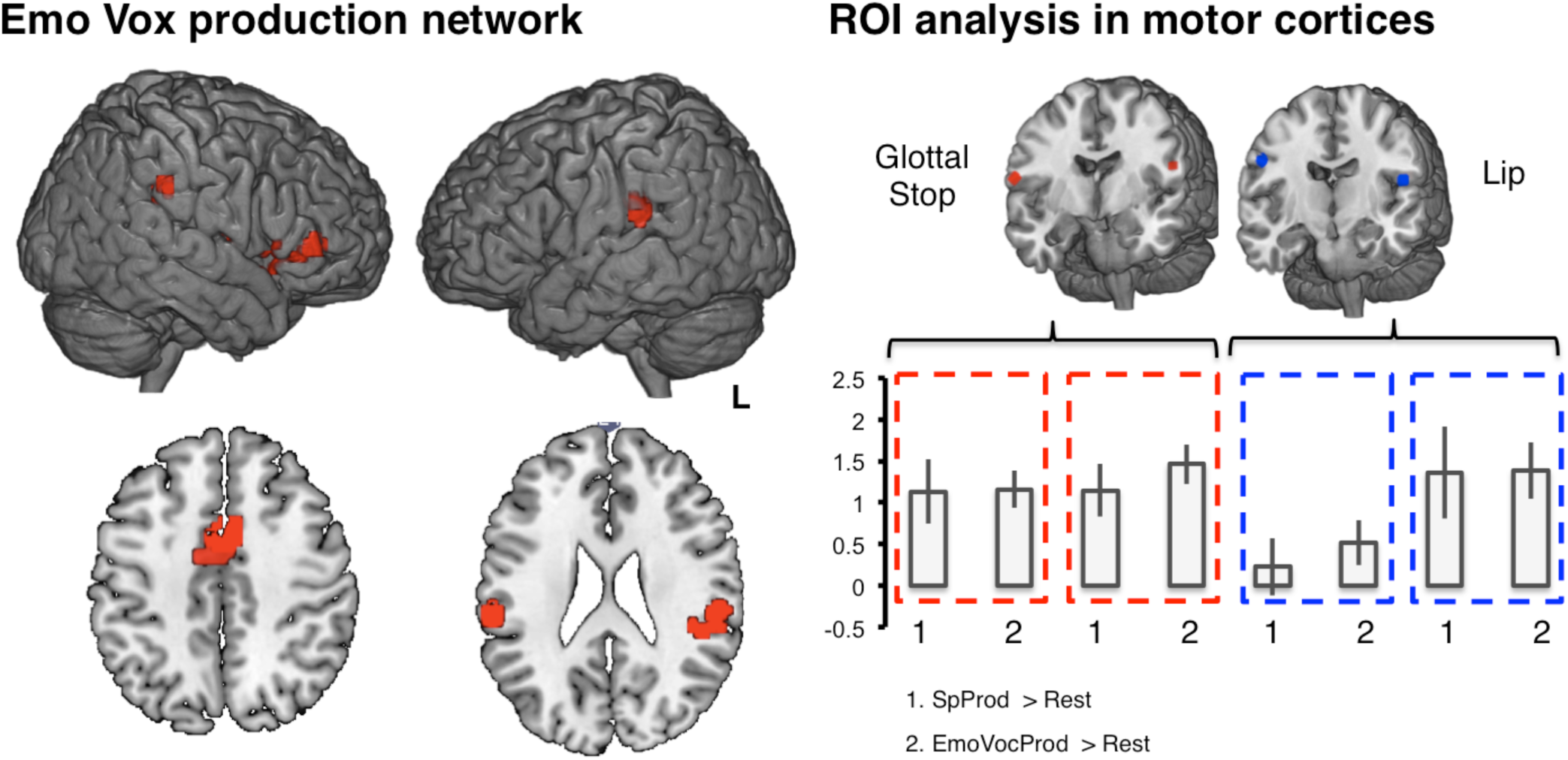
Production of emotional vocalizations compared to speech production is associated with increased activity in widespread cortical regions but not primary sensorimotor cortices. The comparison of production of the two classes of vocalization (Speech and Emotional Vocalizations) revealed significant differences in a network of sensorimotor regions lying outside of primary cortices (EmoVox > Speech production, left panel). These include the putamen (not shown), right cerebellum, bilateral inferior parietal cortex, multiple clusters lying in the left IFG (pars orbitalis, BA47 and pars opercularis, BA44 with a subpeak in pars triangularis, BA45), left insula cortex, right pre-supplementary motor area (premotor cortex), middle temporal gyrus and left medial superior parietal cortex (cuneus, not shown). A region of interest analysis confirmed that BOLD responses did not differ between these two classes of movement in primary motor or somatosensory regions associated with articulatory movements of the upper and lower vocal tract (right panel).

## Discussion

The present study aimed to delineate the neural networks underlying two types of articulatory movements, human speech and emotional vocalizations. We report for the first time that there are clear neural differences underlying the production of these movements on three levels of the motor hierarchy: the internal representation, the way in which afferent information is processed and finally, in terms of execution of these movements. These data provide the first functional neuroimaging data to suggest that these vocal behaviors may comprise two distinct classes of movements, due to a partial dissociation in the neural control of speech and emotional vocal behaviors. First, we show that activity in primary motor and somatosensory cortices are not distinguishable for the production of speech and emotional vocalizations, but non-primary cortical regions do show differential responses. Then, we report patterns of activity during production and miming which indicate that auditory information is processed differently during speech and emotional vocalization production.

Since the advent of positron emission tomography (PET) imaging, the widespread neural networks involved in the production of speech have been well described, including ventral sensorimotor cortices, supplementary motor area, inferior frontal gyri extending to insula cortex, mid to posterior superior temporal gyri and inferior parietal cortices in both hemispheres (for a review, see Price, 2012). In the last decade of so, attention has focused on attempting to figure out the functions of the many nodes in this network. Post-mortem studies have identified direct links between precentral gyrus and the *nucleus ambiguus* of the brainstem (Kuypers, 1958b). Evidence suggests that these connections may be present in chimpanzees but not in squirrel monkeys or cats (Jurgens, 1976; Kuypers, 1958a). It has been proposed that these direct connections between the pre-central gyrus and brainstem may have evolved in higher primates to support, at least in part, laryngeal motor control for human speech (Larson, 1988) although care must be taken to extrapolate from primates who’s vocal repertoire is very different to that of humans. Despite the prominent role of emotional vocalization such as laughter in social communication (Scott et al, TICS 2014), the neural basis of emotional vocalization production is hugely under investigated (Rohrer et al., 2009). In lieu of studying human emotional vocalizations, research has focused on the neural control of non-human primate vocalizations. Based on this and work in humans, Simonyan and Horwitz (Simonyan & Horwitz, 2011) have proposed a three tier system for the motor control of human vocalization. The lowest level comprises the brain stem and phonatory nuclei within the spinal cord that coordinates the control of innate productions via articulatory, laryngeal and respiratory movements. At the midlevel, there is the periaqueductal grey (PAG) anterior cingulate cortex and limbic system, including the thalamus, which is central to the initiation and control of voluntary emotional vocalizations such as those described in the present experiment. Finally at the highest level there is the output system comprised of primary motor cortex corresponding to the laryngeal and orofacial motor regions. These cortical structures are responsible of voluntary production and control of speech and song.

We report a network of regions more active during the production of emotional vocalizations in non-primary fronto-parietal sensorimotor regions, areas that have established roles in primate and non-human primate vocalization. Here we show that these same regions underlie the production of emotional vocalizations, more than for the highly over learnt motor act of speech. These data support clinical findings, demonstrating that patients who are unable to produce speech, can sometimes still produce emotional vocalizations (Dronkers, 1996). For example, Roher and colleagues (Rohrer et al., 2009) describe ten patients with primary progressive aphasia from fronto-temporal damage resulting in mutism in whom their laughter like vocalizations ‘occurred as an automatic vocal output that replaced speech’. It is possible then to imagine that the networks underlying emotional vocalization production might constitute potential therapeutic targets for improving communication in patients with mutism/severe speech production deficits. We suggest that the production of emotional vocalizations, relies on distinct motor processes that may remain intact after damage resulting in loss of speech. If this were true, this second pathway to vocal behavior may constitute a potential alternative therapeutic target for speech rehabilitation.

One subcortical structure which showed a differential response during the production of emotional vocalizations, was the putamen. The putamen comprises the largest output structure from the laryngeal motor cortex. Lesions to this area do not affect monkey vocalizations, but does result in dysarthria and dysphonia in humans (Jurgens, 2002). This is taken as evidence that the putamen plays a central role in learnt voluntary articulations but not innate vocalizations (Simonyan & Horwitz, 2011). On the other hand, Simonyan & Horwitz also suggest that in humans, primary laryngeal and orofacial motor regions are the regions that underlie voluntary production and control of speech and song, as opposed to emotional vocalizations which they suggest should be under the control of the periaqueductal grey (PAG), anterior cingulate cortex and limbic system, including the thalamus. Our finding that the putamen is more active in the voluntary production of emotional vocalizations, a category of innate movements, support this assertion. It remains to be seen how the neural control of naturally occurring emotional vocalizations, compares to that of cued, voluntary emotional vocalization production as studied here.

In addition to this, we report activity in the cerebellum during emotional vocalization production compared to speech production. The cerebellum is a fundamental component of the motor output system involved in integrating afferent information that occurs as a consequence of movement, and using this information, as well as internal models of action, for the coordination and fine sensorimotor control of movement. Accordingly, cerebellar damage results in deficits to speech, most commonly seen as cerebellar dysarthria. Jurgens (2002) asserts that the cerebellum is not necessary for non-speech vocalizations. In squirrel monkeys, lesions to the cerebellar nuclei have been shown not to affect the acoustics of vocalizations (Kirzinger, 1985). However in mice, there is much evidence to suggest a role for the cerebellum in vocalization. Mouse pups lacking the FOXP2 gene, a gene related to speech production, produce dramatically less ultrasonic vocalizations and that this is associated with disrupted development of the cerebellum. Spectral analyses showed however that whilst the incidence of vocalizations was reduced, the acoustic properties of the knockout mice were indistinguishable from the wild type mice (Shu et al., 2005). This suggests a potential role for the cerebellum in the initiation of vocalization; perhaps in addition to the role of feedback processing that is commonly assigned to the cerebellum in motor control (Miall, Reckess, & Imamizu, 2001). In humans also, cerebellar damage is known to affect speech (Gordon, 1996) resulting in dysarthria. Our data suggest that the feedback processing role that the cerebellum plays in the monitoring of fast movements may be distinct for speech and emotional vocalizations.

In the cerebral cortices, we found increased activity in left inferior frontal gyrus, (BA 44 and 47), insula cortex, and OP1 of secondary somatosensory cortex. Left inferior frontal gyrus has a well-established role in the production of articulatory movements, as well as complex hand movements. Bilateral removal of the homologue of Broca’s area in non human primates did not affect performance of trained phonation (Sutton, Larson, & Lindeman, 1974), however in humans lesion studies have shown that loss of the left insula is the greatest predictor of speech production deficits in patients following stroke (Dronkers, 1996). Here we show that this same area is also involved in the production of emotional vocalizations, not just speech. OP1 of secondary somatosensory cortex, left medial superior parietal cortex, marginal gyrus are all part of the multisensory feedback systems that support feedback control during fast vocal movements. The precise role of the SMG in vocalization is unclear. It constitutes a site of multisensory convergence, receiving auditory, somatosensory and visual input, and evidence suggests that in relation to vocal behaviors, this region plays a role in error detection (Shum, Shiller, Baum, & Gracco, 2011), phonological processing (Hartwigsen et al., 2010), and verbal working memory (Deschamps, Baum, & Gracco, 2014). Here we report that producing emotional vocalizations compared to speech elicits activity in the most ventro-rostral aspect of the supramarginal gyrus where it borders with secondary somatosensory cortex.

Lastly, we saw activity in pre SMA during the production of emotional vocalizations. PreSMA, lying between the prefrontal cognitive and precentral motor areas of the frontal lobe, has a well-established role in voluntary decisions to move. Direct stimulation of preSMA results in a desire to move and eventually the execution of that movement (Fried et al., 1991; Penfield & Welch, 1951). A case study describing lesioned pre-SMA but preserved SMA provided compelling evidence that pre-SMA is specifically involved in the inhibition of movement plans especially in the face of other competing motor responses (Nachev, Wydell, O'Neill, Husain, & Kennard, 2007). It is possible that voluntary production of emotional vocalizations drew more heavily on this action selection system, as people are not used to performing these movements on cue, in the way that they may be with regards to speech. Pre-SMA has also been implicated in the preparation of entire sequences of complex movements (Tanji & Shima, 1994).

### The role of recurrent auditory information in different vocal utterances

The present data demonstrate that production of both speech and emotional vocalizations compared to silent miming results in activity in bilateral STG lying close to Heschl’s gyrus (primary auditory cortex). This is unsurprising as the main difference between production and mouthing is the presence of auditory consequences of ones vocal movements. Arguably then, the response of auditory areas in the comparison of production with mouthing, informs us about how the brain is dealing with the afferent information produced as a consequence of making a movement. Outside of this mid STG region, the patterns of activity associated with making a speech sound and making an emotional vocalization look distinct. The sound of self-made speech activates posterior STG and STS whereas the sounds produced as a consequence of making an emotional vocalization is processed in more medial STG regions and the inferior frontal gyrus. While it is hard to say exactly what is happening in the IFG, this does indicate that the way in which sensory and speech responsive cortices process the self-made sounds of speech and emotional vocalizations is fundamentally different. Speech, a highly over learnt motor act, is influenced by modulating recurrent auditory information. For example, introducing time delays in auditory information is highly disruptive to speech production (Howell & Powell, 1987) and perturbing pitch during voicing leads to a rapid compensatory effect in the order of milliseconds (Burnett, Freedland, Larson, & Hain, 1998; Burnett & Larson, 2002). We propose that emotional vocalizations, lying towards the more innate end of the spectrum of complex human movements, may not rely on active feedback monitoring in the manner in which speech does. It would be of interest for future studies to investigate if this extends to emotional facial expressions.

It is well documented that activity in regions of STG are suppressed during self-vocalization. Eliades and Wang (2003, Eliades and Wang 2008) first reported the existence of populations of cells in STG in whom activity in suppressed below baseline levels of activity, during self made vocalizations in macaques. The fact that this suppression begins before the onset of vocalization is evidence that early motor processes drive suppression (Eliades & Wang, 2005, Eliades 2008). They also report populations of cells in neighboring regions that are more activated during self-vocalization, indicating that distinct populations of cells in STG play different roles in processing the auditory consequences of vocal behaviour. In humans this effect, known as speech induced suppression (SIS), has been shown via various modalities (Grix & Watkins, 2010; Harper, Robbins, & Gates, 2010; Zhang, Biggs, & Watkins, 2010) and the effect appears not to be specific to speech (Blakemore, Wolpert, & Frith, 2000; Blakemore, Wolpert, & Frith, 1998; Grix & Watkins, 2010). It is worth noting that some studies have defined speech induced suppression as a reduction in activity during production compared to passive listening, as opposed to a silent baseline condition. The present study did not have a passive listening condition, only an auditory localizer run, so is unable to speak to this type of suppression directly. Here we show differences in how sounds made as a result of speaking or producing emotional vocalizations are processing in auditory regions. It is possible that the extent to which, and the regions where suppression takes place differs for different articulations.

### Neural control of posed/cued vocalizations and naturally occurring vocalizations

Much of the non-human primate literature discussed here pertains to distinct neural systems for the neural control of naturally occurring vocalizations, as opposed to cued or voluntary vocalizations. There is a wealth of data showing that the production of endogenously and exogenously initiated actions in general, involves different neural networks. In contrast, patients with both left and right brain damage have been shown to be equally good at producing posed facial expressions (Caltagirone et al., 1989), and perhaps more interesting was the finding that even patients with oral apraxia were able to produce posed facial expressions. The authors suggest that two distinct neural pathways underlie the neural control of smiling, one being responsible for voluntary expressions and the other with spontaneous smiling. Studies in both humans and non-human primates have revealed that medial supplementary motor areas SMA and lateral pre-motor areas PMA have distinct roles in the production of internally initiated and externally cued movements. More specifically, supplementary motor area plays a relatively more central role in internally driven movements, and pre SMA in more active during the production of externally cued movements (Holt, Kumar, & Watkins, 2010; Johnston, Taliaferro, & Leigh, 2010; Kitagawa et al., 2010). In fact we do report increased activity in pre-SMA in the production of emotional vocalizations compared to speech, which as previously mentioned may relate to the unnatural nature of producing externally cued emotional vocalizations. Obbi and Haggard (Obhi & Haggard, 2004) among many others have shown that internally generated and externally cued vocalizations appear to be controlled by distinct neural structures. With this in mind, we believe it would be of great interest for future work to address the neural basis of voluntary and naturally occurring emotional vocalizations.

## Conclusion

The present experiment set out to look for evidence of a neural dissociation between these two classes of movements, as would be predicted from the non-human primate and human clinical literature. In sum, we have shown that there is partial dissociation between the neural structures underlying the production of human speech and emotional vocalizations, suggesting that the production of emotional vocalizations, may rely on motor processes that may remain intact after damage resulting in loss of speech. This supports clinical findings, asserting that patients with disturbed or lost speech, can have been shown to retain the motor control that elicits normal emotional vocalizations. We report a network of regions, commonly observed in the voluntary control of movement that is more active during cued production of emotional vocalizations compared to speech. Many related functions have been attributed to these cortical and subcortical sites, many of which fit with the constraints of the present task demands. From these data it is not possible to conclude what the specific roles of these different nodes are in the production of emotional vocalizations compared to speech. However these findings confirm significant and widespread differences in the overt production and silent miming of emotional vocalizations, which may purport to preparation, execution, feedback control or feedback processing of these articulatory movements. Future experiments should aim to investigate these different stages of motor processing more closely. We tentatively suggest that the role of auditory feedback may differ across these two classes of vocal movements and propose that further investigation of such questions may point us to novel therapeutic targets for the rehabilitation of speech production disorders.

